# How past exposure to predation affects personality and behavioural plasticity?

**DOI:** 10.1101/844258

**Authors:** Juliette Tariel, Sandrine Plénet, Émilien Luquet

## Abstract

When behaviour is observed multiple times on animals from the same population, between-individual difference in mean behaviour (personality) and in behavioural plasticity are often reported. While the developmental environment might explain such an individual difference, the effect of parental environment is still unexplored -a surprising observation as parental carry-over effect are now well-known to influence the average phenotype and plastic responses of offspring-.

We tested whether parental and developmental exposure to predator cues impacted the immediate behavioural predator-induced response in the snail *Physa acuta* at both mean level (average response) and individual level (between-individual variation). Two generations of snails were reared in laboratory without or with exposure to predator cues. Then, escape behaviour was repeatedly assessed on adult snails in the presence or absence of predator cues.

Both parental and developmental exposure to predator cues acted additively towards a lower average behavioural plasticity. At the individual level, developmental exposure induced higher differentiation in personality trait but not in behavioural plasticity while parental environment did not influence the between-individual differences.

These results highlight that past environments can strongly influence behaviour at the population level and that they should be taken into consideration when investigating why individuals differ in behaviour.

## Introduction

Face to rapid environmental changes, organisms render alternative phenotypes within a generation (phenotypic plasticity) (West-Eberhard 2003; Pigliucci 2005). Plasticity may also occur across generations, when the phenotype of a new generation is influenced by the environment experienced by the previous generation(s) (Agrawal et al. 1999; Galloway and Etterson 2007; Salinas et al. 2013; Donelson et al. 2018). Among plastic traits, behavioural traits are considered as the most flexible and reversible traits, providing the fastest responses to immediate environment (Hazlett 1995; West-Eberhard 2003; Relyea 2005; Ghalambor et al. 2010). It is then predicted that behavioural traits would be less influenced by past environmental experience than by immediate environment (Piersma and Drent 2003; Dingemanse and Wolf 2013). However, developmental conditions were found to influence the average response across many individuals from the same population (West-Eberhard 2003; Stamps and Groothuis 2010). Moreover, few evolutionary biology studies demonstrated that ancestral environments can influence the behaviour of offspring (Bestion et al. 2014; Mateo 2014; Tariel et al. 2019). For instance, juvenile common lizards *Zootoca vivipara* born from mothers exposed to predator cues increased their activity on average when face to predator cues compared to juveniles from non-exposed mothers that decreased their activity (Bestion et al. 2014). It means that past environmental conditions both through development and across generations can constraint highly flexible and reversible average responses to immediate environments.

Developmental environmental conditions are also known to impact personality (Biro et al. 2010; DiRienzo et al. 2015; Urszán et al. 2015; Han and Dingemanse 2017; Royauté and Dochtermann 2017), *i.e.* consistent between-individual difference in mean behaviour across time within a single group (*e.g.* population). For instance, a bacterial injection during development of field crickets *Gryllus integer* induced a lower between-individual variation in boldness, meaning more similar individual behaviour (DiRienzo et al. 2015). In contrast, relatively few studies have shown that developmental environments can affect between-individual variation in plasticity (Dingemanse et al. 2012; Briffa et al. 2013; Gribben et al. 2013; Urszán et al. 2018), *i.e.* individual responsiveness to environmental variation. For instance, exposure to predator cues during the development of *Rana dalmatina* tadpoles induced a higher diversity in individual boldness plasticity to immediate predation risk (Urszán et al. 2018). Personality and between-individual variation in behavioural plasticity has been ignored for a long time, considered as a non-significant noise (Dingemanse and Wolf 2013; Mathot and Dingemanse 2014). In the last decade, the statistical and biological significance of such variation has become widely accepted and studying the causes and consequences of this variation has become a central topic of evolutionary behavioural ecology (Mathot et al. 2011; Giudice 2015). In addition to the developmental environment, we hypothesized that carry-over effects of past environments experienced by the previous generation(s) could also generate between-individual variation of behavioural responses. Indeed, generational carry-over effects are known to influence the average behaviour of offspring and theoretical studies suggest that parental environment could impact how individuals react to immediate environmental conditions (Reddon 2012; Snell-Rood 2013; Giudice 2015; Beaman et al. 2016; Stamps 2016; Rittschof and Hughes 2018) but, to our knowledge, this has never been investigated.

Using the theoretical framework developed by Dingemanse et al. (2010), we investigated how developmental environment experienced by parents (parental environment) and by offspring (developmental environment) influenced average behavioural response and between-individual variation in behavioural response. Based on the concept of behavioural reaction norm (*i.e.* the pattern of behavioural phenotypes that a single individual produces in a given set of environments), this approach allows to combine the study of animal personality and individual plasticity, considering each individual by an elevation/intercept (*personality*, *i.e.* the mean level of behaviour in the average environment) and by a slope (*behavioural plasticity*, *i.e.* the responsiveness to the immediate environment).

Predation is a major selective pressure and has a strong impact on prey behaviour (Lima and Dill 1990). Immediate anti-predator behaviour (*e.g.* escape, refuge use, freezing, vigilance) may be expressed when predator cues are detected (Tollrian and Harvell 1999). Such detection can trigger long-lasting effects on average anti-predator behaviour of future life stages (Relyea 2003; Beaty et al. 2016) and even across generations (Storm and Lima 2010; Giesing et al. 2011; Keiser and Mondor 2013; Bestion et al. 2014; Luquet and Tariel 2016; Tariel et al. 2019), allowing a pre-adaptation to predation risk. At the individual level, although one set of studies highlighted that cues of predator presence during development change the between-individual variation in anti-predator behavioural reaction norm (Urszán et al. 2015, 2018), how parental environmental can influence it is still an open question.

The hermaphroditic *Physa* freshwater gastropods has been widely used for studying predator-induced defences (*e.g.* DeWitt 1998; Auld and Relyea 2011; Gustafson et al. 2014). These snail species display in average a crawling-out (escape) behaviour, a thicker shell and a narrower shell and aperture shape when exposed to predator cues during development (Alexander and Covich 1991; DeWitt 1998; Auld and Relyea 2011; Luquet and Tariel 2016). In addition, the *Physa acuta* species has a short life cycle and has already been used for transgenerational studies of plasticity showing an influence of grand-parental and parental exposure to predator cues on the average escape behaviour (Luquet and Tariel 2016; Tariel et al. 2019).

In order to investigate the effects of past environments (both parental and developmental) on anti-predator behavioural reaction norm, we raised two generations of *P. acuta* snails without or with predator cues. We examined individual reaction norms of escape behaviour on adult snails in response to an immediate predator cues exposure. We tested whether parental and/or developmental exposure to predator cues shaped 1 - the average behavioural response to immediate exposure to predator cues and 2 - the between-individual variation in behavioural reaction norm (personality and individual plasticity).

## Material & Methods

### Animal collection and experimental design

We collected adult *P. acuta* snails (F_0_, n = 86) in a population from a lentic backwater of the Rhône river in Lyon, France (45.80° N, 04.92° E) in February 2017. Snails interbreed (hermaphroditic *P. acuta* performs predominantly outcrossing (Henry et al. 2005)) overnight in a 10 L plastic vial filled with dechlorinated tap water. Next, adult F0 snails were isolated and left to lay eggs for 24h in 70 mL plastic vials, and then removed. 24 vials containing one egg capsule each were randomly kept and constituted our 24 F1 maternal families (hereafter called “family”, *i.e.* progeny of one F0 snail). During the whole experiment, all vials were kept in the same experimental room with a 25°C temperature and a 12h/12h photoperiod. Seven days later, hatched snails were fed *ad libitum* with bowled and blended lettuce, with water and food renewed twice a week. 10 days later, each F1 family was separated in two treatments with: 6 siblings put together in the control (C) water (induced snails) and 6 snails put together in the predator-cue (P) water (non-induced snails). Predator cue water was obtained by individually rearing several crayfish (*Orconectes limosus*) in 4 L dechlorinated tap water and by feeding them twice a week with one smashed *P. acuta* adult. A smashed *P.* acuta adult was also added to the 4 L water one hour before using it. Snails were kept in 6-siblings group for 7 days and then isolated in the same rearing conditions for 16 days (F_1_: n = 288 at the beginning of the experiment with 32 dead snails at the end).

The F2 generation was generated by interbreeding overnight 15 random pairs of F_1_ snails from the same treatment but from different families. Among these 30 F_1_ reproducing snails per treatment, 25 (control F_1_ treatment) and 21 (predator-cue F_1_ treatment) snails finally laid within 24h eggs that constituted the F_2_ families. We then followed the same protocol than before. F_2_ snails were kept in isolation longer (46 days instead of 16 days at the F_1_ generation) to reach a sufficient mass (F_2_: n = 552 at the beginning with 40 dead snails at the end).

In summary, the F_1_ generation consisted in two different developmental environments (control C and predator-cue P) and the F2 generation in 4 combinations of parental and developmental environments (CC, CP, PC or PP) (Figure 1).

**Figure 1.**
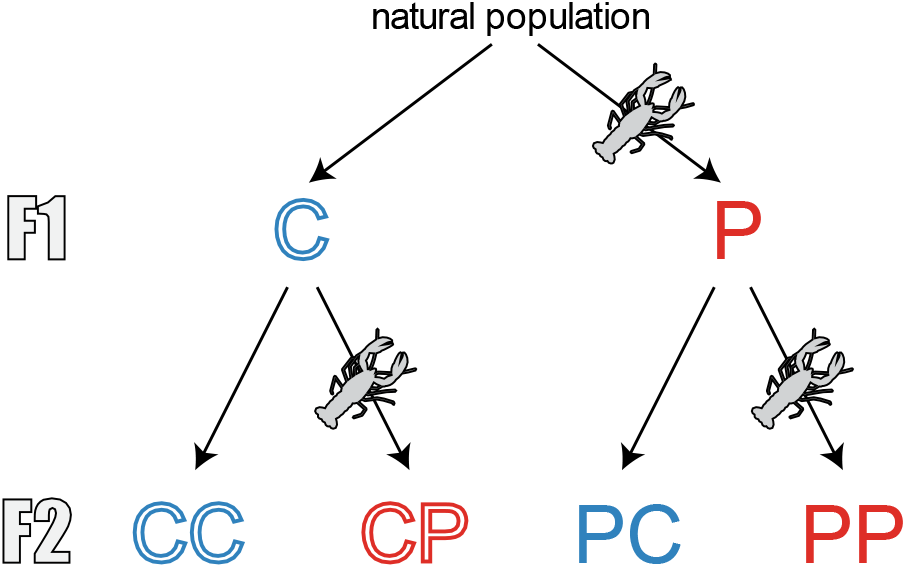
Experimental design at the F1 and F2 generations. The F_1_ generation consisted in two different developmental environments (control C and predator-cue P) and the F2 generation in 4 combinations of parental and developmental environments (CC, CP, PC or PP). For instance, CP illustrated a treatment with a control parental environment and a predator-cue developmental environment. 20 snails were scored for each treatment.

### Behavioural assessments

The escape behaviour of 20 snails per treatment (one snail per family, 20 F1 families and 20 F2 families randomly chosen) was assessed in both immediate control and predator-cue environments, for a total of 40 F1 and 80 F2 snails scored. Escape behaviour was estimated by the time taken by the snail to crawl-out the water, a classic answer to benthic predators like crayfish (Alexander and Covich 1991). It was scored in a rearing vial in which 7 mm polystyrene was placed at the bottom delimiting an acclimation chamber (23 mm diameter) at the centre. Snail was put in the acclimation chamber for 1 min. Then the time to reach the surface was recorded using JWatcher (Blumstein and Daniel 2007) and the experiment stopped after 5 min. The time to crawl-out was scored 4 times for each snail, twice in control water and twice in predator-cue water, to estimate individual reaction norm for each snail. The 4 scores were done within a day, with a maximum of 3h between two consecutive scores. Finally, total mass (body and shell) was measured for each snail with an electronic scale at the nearest 0.0001 g.

### Statistical analysis

The effects of past exposure to predator cues on time to crawl-out (our proxy for escape behaviour) were studied with linear mixed models (LMM). The time to crawl-out was log10-transformed to achieve normality and minus-transformed as a short time to crawl-out is a proxy of a fast escape. F1 et F2 datasets were analysed separately using the same structure of LMM including immediate, developmental and parental (only for the F2 dataset) environments and all interactions (see model equations in supplementary material 1). The main fixed effects were centre prior analysis (contrasts set to −0.5 for “C” control environment and 0.5 for “P” predator-cue environment) allowing to estimate individual intercept values in a theoretical mean environment (Dingemanse et al. 2010). Snail total mass was scaled and added as a fixed covariable to control for size effect. Interactions with mass and trial number were not significant and not included in the fixed effects. The random effect structure depended on the hypothesis tested (see details below). All analyses were done in R 3.4.1 (R Development Core Team 2008).

#### Personality and between-individual variation in plasticity

Based on the fixed effect structure described above, we fitted three models only differing by their random structure to test variance in personality trait and in behavioural plasticity (see model equations in supplementary material 1). LM0, a null model with only a residual variance 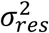. LMM1, a random intercept model with 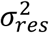 and a variance in individual intercept 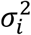. LMM2, a random slope model with 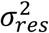, 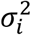, a variance in individual slope to the immediate environment 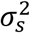 and a correlation between individual intercept and slope 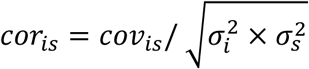. These models were fitted with restricted maximum likelihood estimation using the package lme4 (Bates et al. 2015). We tested the significance of personality (LM0 vs LMM1) and variation in plasticity (LMM1 vs LMM2) using likelihood ratio tests (LRT). Using LMM1, we calculated the repeatability R of the escape behaviour defined as 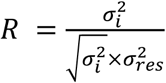.

#### Effects of past environments on personality and between-individual variation in plasticity

Given that we didn’t find any between-individual difference in behavioural plasticity (see results below), we only tested the effects of past environments (developmental and parental) on personality. Based on the fixed effect structure described above, we fitted LMM3, a random intercept model with a residual variance 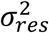 and one variance in intercept 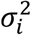 for each treatment (two 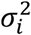 at the F1 and four 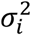 at the F2; see supplementary material 1 for LMM3 equation). We carried out Bayesian Markov chains Monte-Carlo (MCMC) procedures implemented in the MCMCglmm package (Hadfield 2010) to obtain the posterior distributions of parameters, the estimates (mean of posterior distribution) and the 95% confidence intervals (CI). Bayesian p-values were calculated to compare variance in intercept between treatments by dividing the number of iterations fulfilling a condition (for instance, 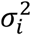 in developmental environment C superior or equal to 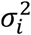 for developmental environment P) by the total number of iterations. The details about the LMM3 modelling procedures is given in supplementary material 1.

#### Effects of past environments on average escape behaviour

To test the effects of past exposure to predator cues on average behavioural response across all individuals, we analysed the fixed effects of the LMM3 described above using the same Bayesian MCMC procedures. With the parameter estimates and the package emmeans (Lenth 2019), we calculated contrasts between estimated means.

## Results

### Effects of past environments on average escape behaviour

At the F1 generation, snails escaped on average 13 sec (12%) faster when exposed to immediate predator-cue (Table 1; Figure 2). The developmental environment did not affect this response (Table 1; Figure 2).

**Table 1.**
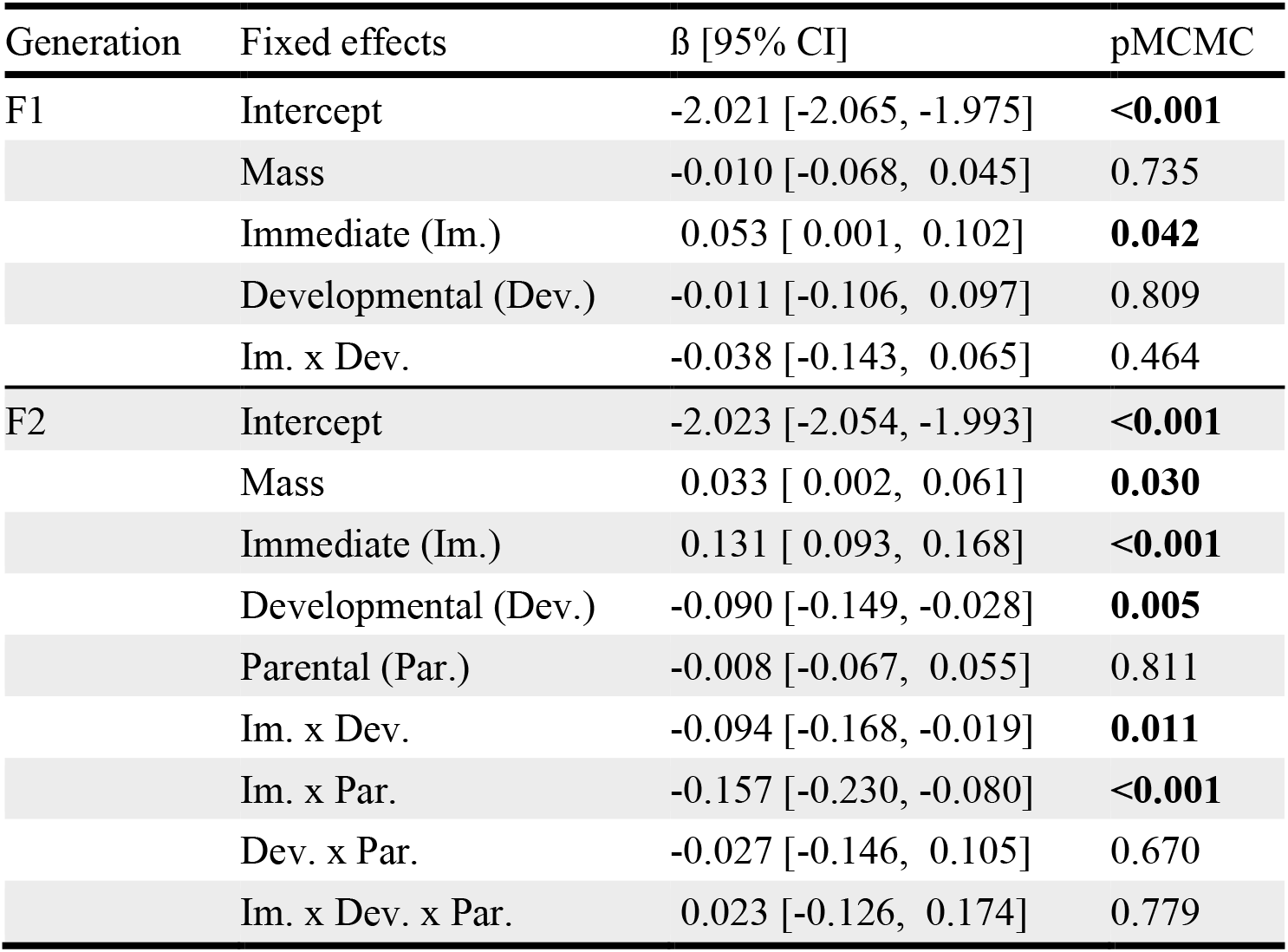
Effects of parental and developmental environments on average escape behavioural response to the immediate environment at the F1 and F2 generations. Parameter estimates (β) of fixed effects are means of the parameter posterior distribution with their 95% confidence interval. pMCMC represents Bayesian p-value and are bold if pMCMC < 0.05. Random effects of this linear mixed model (LMM3) are presented in Table 3. Model equations are available in supplementary material 1.

**Figure 2.**
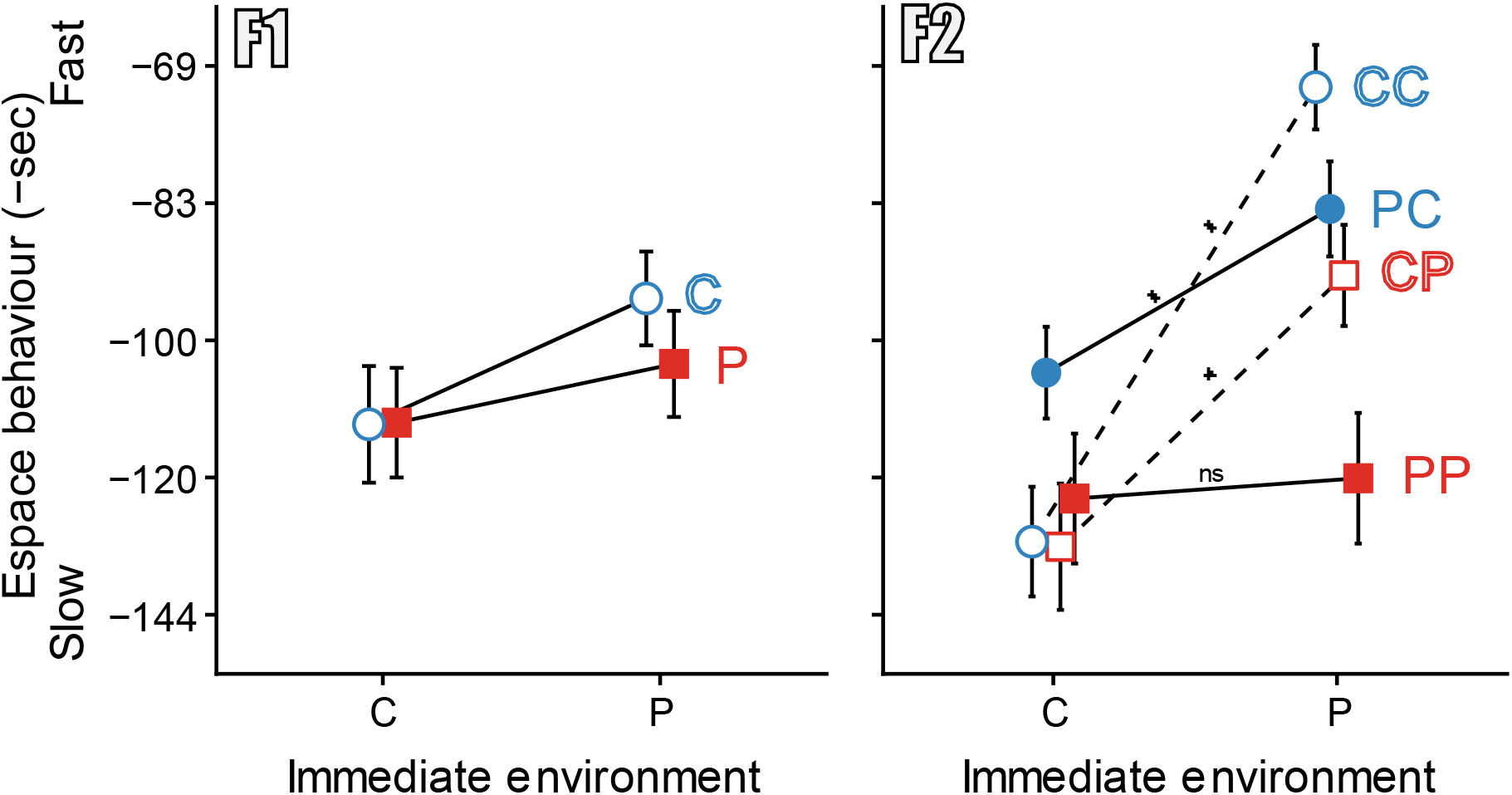
Effects of parental and developmental environments on average escape behavioural response to the immediate environment at the F1 (left panel) and F2 (right panel) generations. The y-axis is minus of log10 of time to crawl-out (a proxy of escape behaviour) with the y-scale back-transformed in minus seconds. The x-axis is the immediate environment with “C” and “P” for control and predator-cue. Developmental C and P environments are represented with blue circle and red square symbols, respectively. Parental C and P environments (F2 generation only) are represented with open symbols/dashed lines and closed symbols/solid lines, respectively. Each combination of past environments is denoted with two letters, for instance “CP” meaning parental C environment and developmental P environment. “*” symbol and “ns” indicate significant and non-significant average reaction norms, respectively. Points are mean ± SE.

At the F2 generation, the average escape behaviour depended on the additive effects of past environments (significant interactions between immediate x developmental and immediate x parental environments; Table 1; Figure 2). In the control immediate environment, nor parental neither developmental environment influenced escape behaviour (Parental environment P vs C: contrast = 0.071 [−0.004, 0.143]; developmental environment P vs C: contrast = −0.043 [−0.115, 0.027]). In the predator-cue immediate environment, both parental and developmental exposure to predator cues induced a slower escape behaviour (20 sec (24%) and 28s (36%) slower, respectively; parental environment P vs C: contrast = −0.086 [−0.156, −0.014]; developmental environment P vs C: contrast = −0.137 [−0.208, −0.061]). This resulted in an absence of average plasticity to immediate environment for PP snails (induced snails from induced parents), *i.e.* flat reaction norm (see supplementary material 1 for pairwise contrasts).

### Personality and between-individual variation in plasticity

Overall, personality was significant at both F_1_ and F_2_ generations (Table 2: comparison LM0 vs LMM1). The repeatability of escape behaviour was 0.32 (95% CI: 0.20-0.61) at the F_1_ generation and 0.28 (95% CI: 0.20-0.45) at the F_2_ generation. However, the behavioural plasticity did not significantly vary between individuals (Table 2: comparison LMM1 vs LMM2).

**Table 2.** Personality and between-individual variation in behavioural plasticity at the F1 and F2 generations. Three models differing by their random structure are tested against each other with likelihood ratio tests (Tested models), whereas the full fixed effects structure was constant and is shown in Table 1. Variance estimates are given with their 95% confidence interval calculated with parametric bootstrap method on 2000 simulations. *LM0* = model null with no random effect (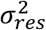, residual variance). *LMM1* = random intercept individual (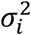, variance in intercept = variation in individual mean behaviour = personality). *LMM2* = random slope by immediate environment (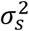, variance in slope = variation in individual behavioural plasticity); cor_is_, correlation between intercept and slope). Bold P-values indicates P < 0.05.

| Generation | Model | $\sigma_{res}^2$ | $\sigma_i^2$ | $\sigma_s^2$ | cor<sub>is</sub> | Tested models | Chisq<sub>df</sub> | P |
| --- | --- | --- | --- | --- | --- | --- | --- | --- |
| F1 | LM0 | 0.038 [0.031, 0.048] |  |  |  |  |  |  |
|  | LMM1 | 0.027 [0.020, 0.033] | 0.012 [0.005, 0.023] |  |  | 0 vs 1 | 16.75<sub>1</sub> | <0.0001 |
|  | LMM2 | 0.025 [0.017, 0.031] | 0.020 [0.008, 0.039] | 0.005 [0.000, 0.024] | -0.862 [-1.000, -0.118] | 1 vs 2 | 3.27<sub>2</sub> | 0.195 |
| F2 | LM0 | 0.040 [0.034, 0.047] |  |  |  |  |  |  |
|  | LMM1 | 0.029 [0.024, 0.034] | 0.011 [0.006, 0.018] |  |  | 0 vs 1 | 26.28<sub>1</sub> | <0.0001 |
|  | LMM2 | 0.028 [0.022, 0.033] | 0.013 [0.006, 0.025] | 0.004 [0.000, 0.018] | -0.392 [-1.000, 1.000] | 1 vs 2 | 0.56<sub>2</sub> | 0.756 |

### Effects of past environments on personality

The developmental exposure to predator cues did not significantly increase personality at the F_1_ generation (Table 3; Figure 3; variance in intercept increased by 2.5-fold; P = 0.147) but significantly increased it by 4-fold at the F_2_ generation (Table 3; Figure 3; P = 0.009). The parental exposure to predator cues increased by 1.3-fold personality but this was not significant (Table 3; Figure 3; P = 0.305).

**Table 3.**
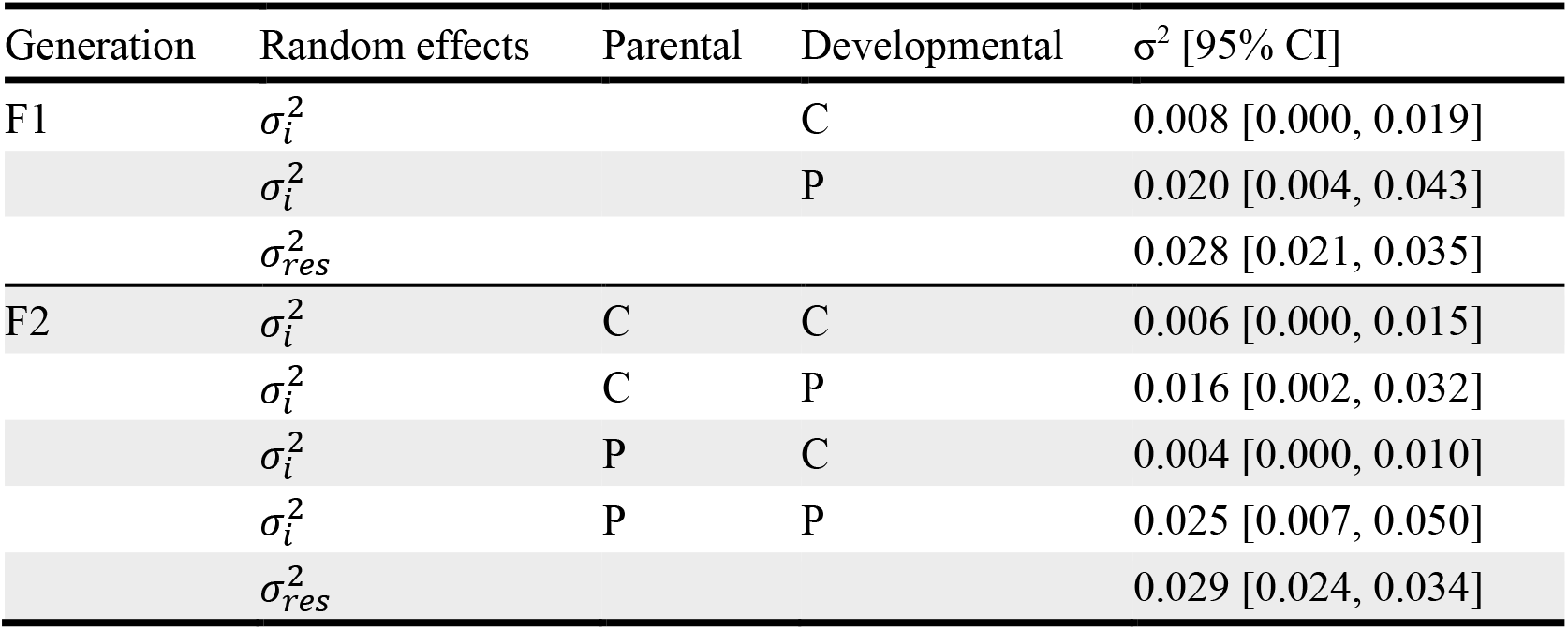
Effects of parental and developmental environments on personality at the F1 and F2 generations. The random part of this model (LMM3) is structured with a random intercept term for each combination of past environments (“C” for control and “P” for predator-cue environments). Mean of variances in intercept 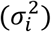 and of residual variances 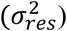 are given with their 95% confidence interval.

**Figure 3.**
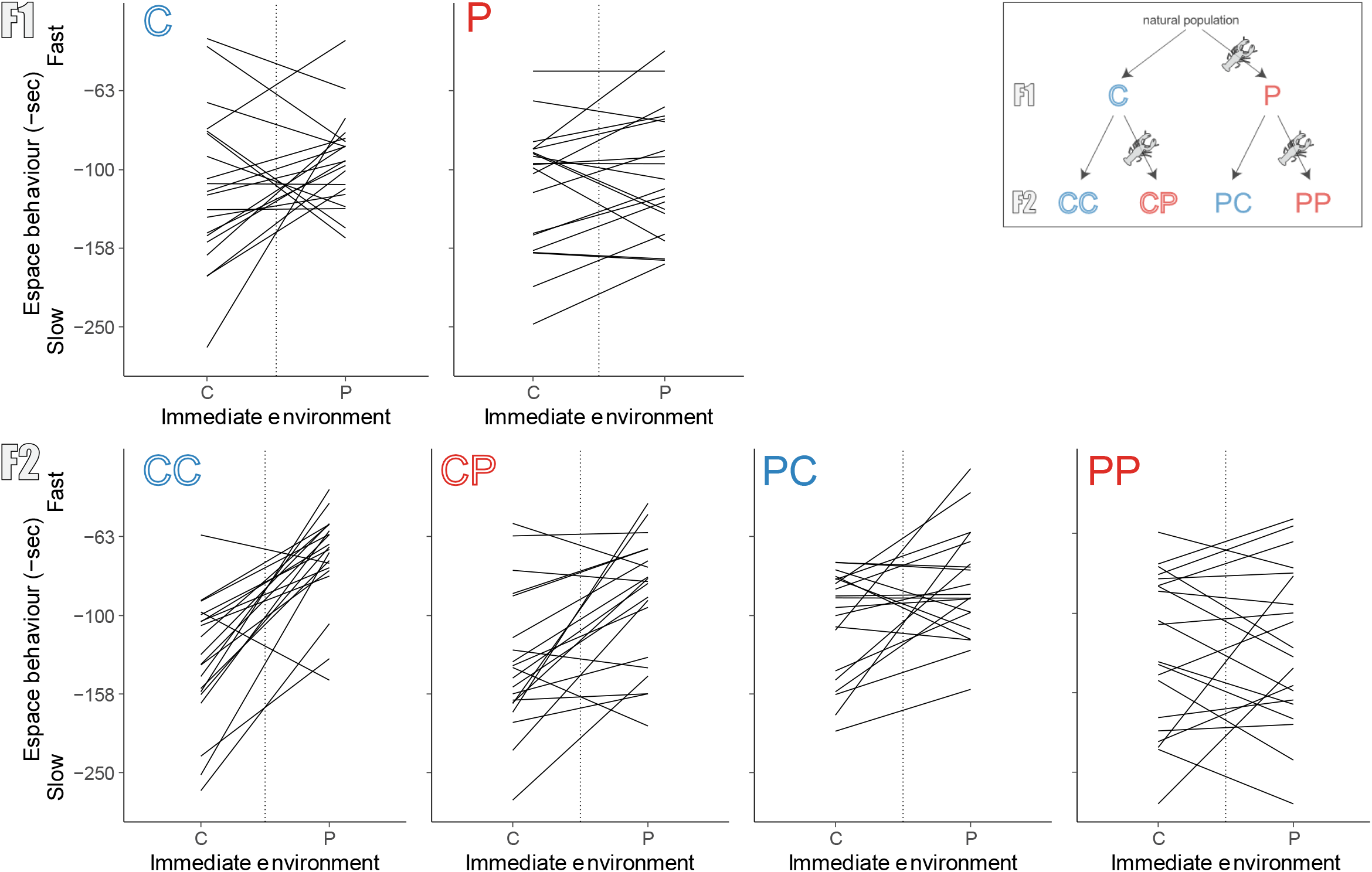
Effects of parental and developmental environments on between-individual variation in escape behavioural response to the immediate environment at the F1 (upper panel) and F2 (bottom panel) generations. The y-axis is minus of log10 of time to crawl-out (escape behaviour) with the y-scale back-transformed in minus seconds. The x-axis is the immediate environment with “C” and “P” for control and predator-cue. On the top-right part of the figure, the Figure 1 is repeated to explain each combination of past environments. The vertical dotted line represents the theoretical average immediate environment where the individual intercept values and then the personality (variance in intercept) are estimated. Each solid line is a reaction norm of one individual, joining the mean of two trials in each immediate environment.

## Discussion

We investigated the effect of both developmental and parental environments on the average behavioural response, the personality and the between-individual variation in behavioural plasticity in the freshwater snail *Physa acuta*. As expected, snails crawled-out the water on average faster facing an immediate exposure to predator (crayfish) cues, the well-known anti-predator behavioural response in *Physa sp.* (Alexander and Covich 1991; DeWitt 1998). Parental and developmental exposure to predator cues additively decreased this average behavioural plasticity to the immediate predation risk at the second generation. We confirmed escape behaviour as a personality trait with consistent between-individual variation in individual mean behaviour. Interestingly, developmental exposure to predator cues resulted in a higher differentiation in personality trait at the second generation while the parental environment did not influence it. However, our study showed that behavioural plasticity did not vary between all individuals.

### Effects of parental and developmental environments on average behaviour

Parental and developmental exposures to predator cues clearly impacted the average behaviour in response to immediate cue exposure at the second generation. The effects of past exposures were context-dependant: parental and developmental environments influenced the escape behaviour only in immediate predator-cue context, inducing a slower escape. Trans- and within-generational carry-over effects of predator exposure might be conflicting cues in a control immediate environment whereas they might give relevant information in a predation-cue context. In the literature, parental experience to predator cues results in most cases in a higher expression of anti-predator behaviour, suggesting a pre-adaptation to predator presence (*e.g.* Storm and Lima 2010; Giesing et al. 2011; Luquet and Tariel 2016). In contrast, a previous experience of predator cues can also reduce the anti-predator behaviour (Donelan and Trussell 2018; Urszán et al. 2018) like in our study. Finally, both developmental and parental exposure to predator cues reduced the anti-predator behavioural plasticity of F_2_ snails, until an absence of response for F2 induced snails (*i.e.* snails exposed to predator cues during their development) from induced parents (PP snails) to immediate predation risk.

Such a pattern could result from a compensation for the anti-predator behaviour by the transgenerational and developmental induction of morphological defences (*e.g.* Agrawal et al. 1999; Bestion et al. 2014; Ahlgren et al. 2015; Luquet and Tariel 2016) compensating for the anti-predator behaviour. Indeed, if morphological defences are (1) less costly to maintain that behavioural defences, (2) slower to produce than behavioural defences, and (3) enough to protect a prey from predation, behavioural defences should work early in the anti-predation sequence before disappearing, leaving time to preys to develop efficient morphological defences. This allows to save the costs of having both morphological and behavioural defences at the same time (Lima and Dill 1990; Chivers et al. 2007; Dijk et al. 2016).

Strong habituation to predator odour could lead to a reduced anti-predator behaviour. Habituation is defined as a simple form of learning, where behaviour response to a persistent stimulus is reduced (Christoffersen 1997). By filtering a constant stimulus, habituation allows an animal to focus its cognitive resources on novel or changing stimulus and better respond to it. Neural mechanisms underlying olfactory habituation are started to be uncovered (Das et al. 2011). Long-term habituation has been highlighted in several animal taxa including gastropods (Christoffersen 1997), with habituation that can persist a while after the disappearance of the stimulus. In our case, snails were submitted for 53 days to the predator odour and behavioural assessments start few days after stopping. Induced snails might then be habituated to this predator odour and might reduce their response to a novel exposure of this same odour. However, transgenerational transmission of habituation has never been highlighted to our knowledge, even if transgenerational transfer of conditioning or sensory imprinting to an odour have already been described in nematods and rodents (Remy 2010; Dias and Ressler 2014).

### Effects of parental and developmental environments on personality

Our study evidenced the crawling-out (escape behaviour) as a personality trait in *P. acuta* with a repeatability estimate around 0.3 at each generation, similarly to other types of behaviour on various animals (Bell et al. 2009). Personality is thought to impact a vast range of ecological and evolutionary dynamics, but our understanding of their proximate causes is still limiting (Stamps and Groothuis 2010; Wolf and Weissing 2012). Here, developmental predation exposure increased the between-individual variance in mean escape behaviour, but the increase was only significant in F_2_ *P. acuta* snails. This result is in accordance with another study (Urszán et al. 2015), suggesting that developmental experience of predator cues generates more variable and extreme anti-predator behaviour. But contrary to our expectations, parental environment did not clearly increase differentiation in personality trait. Parental environment might just impact the average behaviour and not the inter-individual variance, but there is to our knowledge any other empirical studies to compare and generalize this assumption. Moreover, why some past conditions increase or decrease variation among individuals is still unknown. Some studies highlighted, like in our case, an increase of individual differentiation under stressful developmental conditions (predation (Urszán et al. 2015); diet quality (Paaby and Rockman 2014; Royauté and Dochtermann 2017)) while others found the opposite, an increase of variance under favourable developmental conditions (bacterial injection (DiRienzo et al. 2015); diet quality (Han and Dingemanse 2017)). To explain the increase of variance under non-favourable (predation) conditions, we suggest that previous experience to predator cues might trigger more diverse investment strategies between production of anti-predator defences and other traits, such as for example reproduction (*e.g.* earlier reproduction or production of more or higher quality offspring at the expense of production of defences) (Wolf et al. 2007; Urszán et al. 2018). We also suggest that a stressful developmental environment (here predator cues) could increase the level of developmental noise, generating higher variance in personality. Developmental noise is a third source of phenotypic variation, with genetic and environmental effects.

Developmental noise is enough to produce variation in personality under non-stressful conditions. Indeed, clonal animals experiencing nearly identical environments and with no social contact develop consistent differences in personality traits (Vogt et al. 2008; Kain et al. 2012; Bierbach et al. 2017). Level of developmental noise can be modulated by external environments and stressful environments have already been highlighted to increase developmental noise on morphological traits, generating a higher variance in morphology (Willmore et al. 2007; Lazić et al. 2015). However, there is currently to our knowledge any empirical studies showing a causal link between developmental environments and more diverse personality trait through developmental noise.

### Between-individual variation in behavioural plasticity

Despite the recent interest for variation in behavioural plasticity and its consequences on the ecology and evolution of species (Dingemanse et al. 2012; Stamps 2016; Mitchell and Biro 2017), the causes of this variation are unknown. Few studies showed that the developmental environment could modulate between-individual variation in behavioural plasticity (Dingemanse et al. 2012; Briffa et al. 2013; Gribben et al. 2013) as it is the case for personality, but none investigate whether parental environment could also influence variation in behavioural plasticity. Here, we didn’t find any significant between-individual difference in behavioural plasticity at the first and second generations. Individuals were similarly plastic with individual reaction norms close to the average reaction norm of their past environmental treatment. This absence of variation suggests a low genetic diversity in reaction norms in our population. Past predation history over large generational scales might have select and canalize all individuals towards a unique and fixed optimal behavioural plasticity (Kim 2016). However, caution should be taken in overinterpreting absence of between-individual in behavioural plasticity as this variation is hard to detect and requires large data sets (Martin et al. 2011; van de Pol 2012).

### Conclusion

Taken together, our results demonstrated strong impacts of within- and trans-generational environments on both individual and average behavioural responses to predation. A labile and reversible trait like behaviour can be thus constrained by past environments even over generations. While our studies confirmed that developmental environments plays a major role in generating personality, the importance of parental environments on between-individual differences (personality and plasticity) need to be more extensively study.

## Supporting information

Supplementary material 1 - Models equations, Bayesian modelling procedures, pairwise constrasts

## Acknowledgements

We thank E. Desouhant for the careful and critical reading of our manuscript. We would like to thank the following students for their great support with experiments: Laurent Bensoussan, Justine Boutry, Rachel Arnaud, Marie Bouilloud and Anaïs Seve-Minnaert.

## Data accessibility

Data and R code will be moved to an external repository upon publication.

## Authors’ contributions

JT carried out the data analysis and drafted the manuscript; SL designed the study, collected the data, helped draft and critically revised the manuscript; EL designed the study, collected the data, participated in data analysis, helped draft and critically revised the manuscript. All authors gave final approval for publication and agree to be held accountable for the work performed therein.

