## Supplementary material 1 - Models equations, Bayesian modelling procedures, pairwise constrasts for "How past exposure to predation affects personality and behavioural plasticity?"

### Supplementary data from “Parental and developmental exposure to predator cues effects on average escape behaviour, individual personality and behavioural plasticity”

*J. Tariel, S. Plénet & É. Luquet*

*15 novembre, 2019*

#### Contents

**1. Model equations** **1**

**2. Bayesien modelling procedures in MCMCglmm for LMM3** **4**

**3. Contrasts for significance of average reaction norms** **5**

#### 1. Model equations

The effects of past exposure to predator cues on time to crawl-out (our proxy for escape behaviour) were studied with linear mixed models (LMM) and not with a survival analysis as just a few trials were censored (snails that did not crawl-out the water within the 300 sec test; 2 trials out of 160 for the F1 generation; 12 trials out of 320 at the F2 generation; these trials were included in the dataset and set to 300 sec).

In all linear models, $Y_{jk}$ is a single measurement of escape behaviour (time to crawl-out the water in seconds) of individual j (j spreads from 1 to 40 (F1 generation) and from 1 to 80 (F2 generation)) and of trial number k (k spreads from 1 to 4, i.e. four trials for each individual). The residual error of the individual j at the trial k is $E_{jk}$. The residual variance is $\sigma^2_{res}$.

In our paper, fixed effects parameters were estimated with the model 3. They are then only detailed in this model.

**Linear model 0 (null model)**

$-log10(\text{Y}_{jk}) = \beta_0 + \text{fixed effects} + \text{E}_{jk}$

$[\text{E}_{jk}] \overset{i.i.d.}{\sim} \mathcal{N}(0, \sigma^2_{res})$

$\beta_0$ is the parameter for the average intercept.

**Linear mixed model 1 (random intercept model)**

$-log10(\text{Y}_{jk}) = \beta_0 + \text{fixed effects} + \text{U}_{0i} + \text{E}_{jk}$

$[\text{E}_{jk}] \overset{i.i.d.}{\sim} \mathcal{N}(0, \sigma^2_{res})$

$[\text{U}_{0j}] \overset{i.i.d.}{\sim} \mathcal{N}(0, \sigma^2_i)$

The individual intercept value for individual j is $\text{U}_{0j}$ and the between-individual variance in intercept (personality) is $\sigma^2_i$.

**Linear mixed model 2 (random slope model)**

$-log10(\text{Y}_{jk}) = \beta_0 + \text{fixed effects} + \text{U}_{0j} + \text{U}_{1j} \times x^{(I)}_{jk} + \text{E}_{jk}$

$[\text{E}_{jk}] \overset{i.i.d.}{\sim} \mathcal{N}(0, \sigma^2_{res})$

$$\begin{bmatrix} \text{U}_{0j} \\ \text{U}_{1j} \end{bmatrix} \overset{i.i.d.}{\sim} \mathcal{N}(\begin{bmatrix} 0 \\ 0 \end{bmatrix}, \begin{bmatrix} \sigma^2_i & cov_{is} \\ cov_{is} & \sigma^2_s \end{bmatrix})$$

$x^{(I)}_{jk}$ is the immediate environment for the individual j with:

- $x^{(I)}_{jk}$ = -0.5 for k = 1 and k = 2 trial numbers (first two trials in control immediate environment)
- $x^{(I)}_{jk}$ = 0.5 for k = 3 and k = 4 trial numbers (last two trials in predator-cue immediate environment)

The individual slope (behavioural plasticity) value for individual j is $\text{U}_{1j}$ and the between-individual variance in slope is $\sigma^2_s$. The covariance between intercept and slope is $cov_{is}$ and it uses for the calculation of the correlation between intercept and slope: $cor_{is} = \frac{cov_{is}}{\sqrt{\sigma^2_i \times \sigma^2_s}}$.

**Linear mixed model 3 (model with random intercepts for each combination of past environments)**

**F1**

$-log10(\mathrm{Y}_{jk}) = \beta_0 + \beta_1 \times x_j^{(W)} + \beta_2 \times x_{jk}^{(I)} + \beta_3 \times x_j^{(D)} + \gamma_1 \times x_{jk}^{(I)} \times x_j^{(D)} + \mathrm{U_0}_i^{\mathrm{C}} \times \mathrm{I^C} + \mathrm{U_0}_i^{\mathrm{P}} \times \mathrm{I^P} + \mathrm{E}_{jk}$

$[\mathrm{E}_{jk}] \overset{i.i.d.}{\sim} \mathcal{N}(0, \sigma_{res}^2)$

$[\mathrm{U_0}_i^{\mathrm{C}}] \overset{i.i.d.}{\sim} \mathcal{N}(0, \sigma_i^{2,\mathrm{C}})$

$[\mathrm{U_0}_i^{\mathrm{P}}] \overset{i.i.d.}{\sim} \mathcal{N}(0, \sigma_i^{2,\mathrm{P}})$

<u>Fixed effects:</u>

$\beta$ are the parameters for the main fixed effects and $\gamma$ is the parameter for the double interaction term.

$x_j^{(W)}$ is the snail total weight for the individual j.

$x_j^{(D)}$ is the developmental environment for the individual j with:

- $x_j^{(D)}$ = -0.5 for non-induced snails
- $x_j^{(D)}$ = 0.5 for predator-induced snails

<u>Random effects:</u>

For non-induced snails, the individual individual intercept value is $\mathrm{U_0}_i^{\mathrm{C}}$ and the between-individual variance in intercept (personality) is $\sigma_i^{2,\mathrm{C}}$. The same logic applied for predator-induced snails.

$\mathrm{I^C}$ and $\mathrm{I^P}$ are dummy variables:

- $\mathrm{I^C} = 1$ if individual j was not induced, 0 otherwise
- $\mathrm{I^P} = 1$ if individual j was induced, 0 otherwise

**F2**

$-log10(\mathrm{Y}_{jk}) = \beta_0 + \beta_1 \times x_j^{(W)} + \beta_2 \times x_{jk}^{(I)} + \beta_3 \times x_j^{(D)} + \beta_4 \times x_j^{(P)} + \gamma_1 \times x_{jk}^{(I)} \times x_j^{(D)} + \gamma_2 \times x_{jk}^{(I)} \times x_j^{(P)} + \gamma_3 \times x_j^{(D)} \times x_j^{(P)} + \delta_1 \times x_{jk}^{(I)} \times x_j^{(D)} \times x_j^{(P)} + \mathrm{U_0}_i^{\mathrm{CC}} \times \mathrm{I^{CC}} + \mathrm{U_0}_i^{\mathrm{CP}} \times \mathrm{I^{CP}} + \mathrm{U_0}_i^{\mathrm{PC}} \times \mathrm{I^{PC}} + \mathrm{U_0}_i^{\mathrm{PP}} \times \mathrm{I^{PP}} + \mathrm{E}_{jk}$

$[\mathrm{E}_{jk}] \overset{i.i.d.}{\sim} \mathcal{N}(0, \sigma_{res}^2)$

$[\mathrm{U_0}_i^{\mathrm{CC}}] \overset{i.i.d.}{\sim} \mathcal{N}(0, \sigma_i^{2,\mathrm{CC}})$

$[\mathrm{U_0}_i^{\mathrm{CP}}] \overset{i.i.d.}{\sim} \mathcal{N}(0, \sigma_i^{2,\mathrm{CP}})$

$[\mathrm{U_0}_i^{\mathrm{PC}}] \overset{i.i.d.}{\sim} \mathcal{N}(0, \sigma_i^{2,\mathrm{PC}})$

$[\mathrm{U_0}_i^{\mathrm{PP}}] \overset{i.i.d.}{\sim} \mathcal{N}(0, \sigma_i^{2,\mathrm{PP}})$

<u>Fixed effects:</u>

$\delta$ is the parameter for the triple interaction term.

$x_j^{(P)}$ is the parental environment for the individual j with:

- $x_j^{(P)}$ = -0.5 for snails with non-induced parents
- $x_j^{(P)}$ = +0.5 for snails with predator-induced parents

Random effects:

For non-induced snails from induced parents, the individual individual intercept value is $U_{0i}^{PC}$ and the between-individual variance in intercept (personality) is $\sigma_i^{2,PC}$. The same logic applied for CC, CP and PP snails.

$I^{CC}$, $I^{CP}$, $I^{PC}$, $I^{PP}$ are dummy variables:

- $I^{CC}$ = 1 if individual j was not induced from non-induced parents, 0 otherwise
- $I^{CP}$ = 1 if individual j was induced from non-induced parents, 0 otherwise
- $I^{PC}$ = 1 if individual j was not induced from induced parents, 0 otherwise
- $I^{PP}$ = 1 if individual j was induced from induced parents, 0 otherwise

---

#### 2. Bayesien modelling procedures in MCMCglmm for LMM3

We used a non-informative prior with an inverse-Wishart distribution (expected variance V = 1; degree of belief $\nu$ = 0.002; corresponding to a Gamma distribution with shape = 1 and scale = 1). We used a chain of 700 000 iterations with a thin of 100 iterations (1 out of 100 iterations kept) and a burnin of 200 000 iterations (the first 200 000 iterations were discarded). The posterior distribution of each fixed effects parameters and random variances were then made on the 5 000 remaining iterations after thinning and burning. The convergence of MCMC algorithms was checked by: (i) running separately three other chains and analysing the between-chain variance with Gelman & Rubin diagnostic implemented in the package coda (Plummer et al. 2006); (ii) looking at the trace and shape of the posterior distribution; (iii) looking at the values of 95 CI at each iteration and for each parameter; and (iv) calculating the effective size of our parameters (5 000 independent iterations for variances and at least 4 500 independent iterations for fixed effects parameters). The sensibility to prior was also checked by: (i) drawing prior and posterior distribution on the same graph; and (ii) ran two other chains with different priors (inverse-Wishart with V = 0 and $\nu$ = -2; inverse-Wishart with V = 1 and $\nu$ = 0).

---

### 3. Contrasts for significance of average reaction norms

We calculated pairwise contrasts between estimated mean (of LMM3) in immediate environment P vs estimated mean in immediate environment C for each combination of past environments and for each generation. This method allows to assess the significance of average reaction norm (slope of reaction norm different from zero if 95% CI did not overlap zero). Mean and 95% CI of contrasts were calculated with the package emmeans.

| Generation | Environmental history | Contrast immediate environment P vs C |
|---|---|---|
| F1 | C | 0.072 (-0.004, 0.144) |
| | P | 0.034 (-0.036, 0.107) |
| F2 | CC | 0.262 ( 0.185, 0.336) |
| | CP | 0.157 ( 0.082, 0.229) |
| | PC | 0.094 ( 0.021, 0.169) |
| | PP | 0.011 (-0.062, 0.088) |

We found no significant average reactions norms at the F1 generation. We found significant average reactions norms for all environmental histories except PP (induced snails from induced parents) at the F2 generation.
